# Neuronal Hyperactivity Induces Tau Phosphorylation Selectively in Excitatory Pyramidal Neurons Which Contributes to GluA2 Regulation

**DOI:** 10.64898/2026.10.26.762028

**Authors:** Manyi Nina Sun, Kevin Vaughey, Bian Liu, Haidan Guo, Richard Huganir, Meaghan Morris

**Affiliations:** Department of Pathology, Johns Hopkins School of Medicine, Baltimore, MD, 21205; Department of Neuroscience, Johns Hopkins School of Medicine, Baltimore, MD, 21205; Kavli Neuroscience Discovery Institute, Baltimore, MD 21205

## Abstract

While tau plays a role in regulating neuronal network excitability, a process thought to be dysregulated in Alzheimer’s disease (AD), the molecular mechanisms linking tau and hyperexcitability remain unclear. To investigate interactions between endogenous tau and neuronal activity, we increased neuronal activity using bicuculline or suppressed activity using tetrodotoxin for 24 hours in rat hippocampal neurons. Chronic neuronal hyperactivity increased dendritic tau phosphorylation at S202/T205. This response was observed only in excitatory CA1 and CA3 pyramidal neurons, but not in inhibitory or dentate granule neurons. Dendritic phospho-tau positively correlated with neuronal activation after bicuculline treatment, as measured by cfos expression, and was prevented with inhibition of protein kinase A (PKA). To examine the functional consequences of tau S202/T205 phosphorylation, hippocampal neurons were transfected with wild-type tau or tau containing mutations preventing phosphorylation at these sites. Wildtype tau overexpression increased tau phosphorylation and reduced neuronal surface GluA2, which was prevented by phospho-site mutation. Extending these findings to mice, chronic low-dose picrotoxin treatment increased endogenous S202/T205 tau phosphorylation in CA3 neurons. In picrotoxin-treated mice, cfos expression was associated with increased tau phosphorylation specifically in CA3 and CA1 neurons. Together, these findings identify activity-dependent, endogenous tau phosphorylation as a selective response to hyperexcitation in excitatory hippocampal neurons downstream of PKA activation. The data further link tau phosphorylation to altered GluA2 regulation, suggesting a potential molecular connection between neuronal hyperactivity, tau phosphorylation, and synaptic dysfunction relevant to AD.

**Significance Statement:** Neuronal hyperactivity induced endogenous tau phosphorylation at S202/T205 selectively in excitatory hippocampal pyramidal neurons, the same neurons vulnerable to tau aggregation in Alzheimer’s disease (AD). Tau phosphorylation at this site was sufficient to induce downregulation of surface GluA2, implicating phosphorylation of endogenous tau in regulating the response to neuronal hyperactivity. These findings suggest that neuronal hyperactivation modulates tau phosphorylation, potentially linking neuronal hyperexcitation, selective neuronal vulnerability, and synaptic dysfunction in AD.

## Introduction

The microtubule associated protein tau (*MAPT*) plays a role in the regulation of neuronal network excitability, a process thought to be dysregulated in Alzheimer’s disease(Bakker et al., 2012; Vossel et al., 2016; Vossel et al., 2017). Studies of neuronal network excitability consistently show that removal of endogenous tau reduces seizure susceptibility in pharmacologic and genetic models with seizure predisposition(Roberson et al., 2011; Sanchez et al., 2012; DeVos et al., 2013; Holth et al., 2013; Gheyara et al., 2014; Morris et al., 2015b). Subsequent studies have shown that endogenous tau regulates the balance between excitation and inhibition in the brain(Chang et al., 2021). This interaction between tau and neuronal excitability may in part be mediated by regulation of tau phosphorylation. Kainic acid-induced hyperexcitation in mice causes dynamic, time-dependent changes in tau phosphorylation in the brain(Liang et al., 2009). However, the molecular mechanisms linking tau and neuronal hyperexcitation remain unclear.

One potential mechanism linking tau to neuronal network excitability involves regulation of postsynaptic AMPA receptors. Post-synaptic AMPA receptors are the primary mediators of basal synaptic transmission and play a critical role in regulating network homeostasis during chronic changes in neuronal network activity(Fernandes and Carvalho, 2016). Overexpression of tau with mutations inducing aggregation or mimicking high levels of tau phosphorylation induces down-regulation of postsynaptic AMPA receptors(Hoover et al., 2010; Miller et al., 2014), which appears to depend on phosphorylation within the proline-rich domain of tau(Teravskis et al., 2020). Supporting the physiologic role of tau in AMPA receptor regulation, removal of endogenous tau alters the regulation of GluA1 and GluA2 during acute changes in neuronal activity(Suzuki and Kimura, 2017; Prikas et al., 2022). These studies raise the possibility that tau, more specifically changes in tau phosphorylation, may also play a role in chronic neuronal activity changes via AMPA receptor regulation.

To test the role of tau in neuronal hyperexcitability, we took advantage of the homeostatic scaling experimental paradigm. In this paradigm, neuronal activity is chronically increased or decreased without inducing neuronal toxicity. We identified selective changes in tau phosphorylation associated with neuronal activity in cultured neurons and in mice which may contribute to GluA2 down-regulation during chronic neuronal hyperexcitation.

## Materials and Methods

### Primary Neuronal Culture

Primary rat hippocampal culture was performed as previously(Zhang et al., 2021). Briefly, hippocampal neurons were obtained from embryonic day 18 Sprague-Dawley rats. The neurons were plated in standard 12-well tissue culture dishes at a density of 100,000-150,000 cells per well. For tau mutation experiments, neurons were transfected using Lipofectamine 2000 (11668030, Invitrogen, Carlsbad, CA, USA) on DIV 14-15, then fixed on DIV 18-19. For PKA activity experiments, neurons were plated and maintained in standard neuronal media except using Neurobasal medium which lacked phenol red (12348017, Gibco ThermoFisher, Waltham, MA, USA), with transfection at DIV18.

For most experiments, neurons were treated for 24 hours with 1 µM tetrodotoxin citrate (1069, Tocris biosciences, Bristol, UK) or 20 µM bicuculline methiodide (2503, Tocris biosciences, Bristol, UK) for 24 hours starting on DIV18-20. For kinase inhibitor screening, neurons were pre-treated with each inhibitor 10 minutes prior to adding vehicle or bicuculline for 24 hours. Neurons were treated with 2mM lithium chloride (L9650, Sigma-Aldrich, Burlington, MA, USA), 100nM GO6983 (2285, Tocris biosciences, Bristol, UK), 20 µM (S)-Roscotivine (ALX-350-293, Enzo Life Sciences, New York, NY, USA), 20 µM IC261(6676, Tocris biosciences, Bristol, UK), 50 µM TBB(2275, Tocris biosciences, Bristol, UK), 20 µM SB203580 (ab120162, Abcam, Cambridge, UK), 20 µM Ly294002 (1130, Tocris biosciences, Bristol, UK), 10 µM U0126 (1144, Tocris biosciences, Bristol, UK), 5 and 10 µM H89 dihydrochloride (2910, Tocris biosciences, Bristol, UK), or 3 µM KT5720 (1288, Tocris biosciences, Bristol, UK).

### Antibodies and Western Blotting

Primary antibodies and dilutions used in this study were 1:1000 total tau (ab76128, Abcam, Cambridge, UK), 1:1000 phospho-tau S202/T205 (MN1020, ThermoFisher, Waltham, MA, USA), 1:1000 phosphotau T231 (MN1040, ThermoFisher, Waltham, MA, USA), 1:1000 MAP2 (NB300-213, Novus Biologicals, Centennial, CO, USA), 1:400 CTIP2 (AB18465, Abcam, Cambridge, UK), 1:400 NeuroD6 (AB85824, Abcam, Cambridge, UK), 1:150 GAD67 (sc28376, Santa Cruz Biotechnology, Dallas, TX, USA), 1:2000 GFP (ab13970, Abcam, Cambridge, UK), 1:2000 Myc (clone 2235, Huganir lab), 1:1000 GluA2 (purified 15F1, Gouaux lab(Yu et al., 2021)), and 1:500 cfos (226008, Synaptic Systems, Göttingen, Germany). Secondary antibodies were used at a dilution of 1:10,000 for western blot and 1:400 for immunofluorescence. The secondary antibodies used in this study were goat anti-mouse 880CW (926-32210, LI-COR Biosciences, Lincoln, NE, USA), donkey anti-rabbit 680LT (926-68023, LI-COR Biosciences, Lincoln, NE, USA), anti-mouse IgG1 Alexa 488 (A21121, ThermoFisher, Waltham, MA, USA), anti-rabbit Alexa 568 (A11036, ThermoFisher, Waltham, MA, USA), anti-chicken IgY Alexa 405 (A48260, ThermoFisher, Waltham, MA, USA), anti-rat Alexa 647 (A21247, ThermoFisher, Waltham, MA, USA), and anti-mouse IgG3 DyLight 405 (Jackson Immunoresearch Laboratories, West Grove, PA, USA). Western blotting was performed as previously described(Vossel et al., 2015) except for the use of BOLT 4-12% bis-tris plus mini protein gels (NW04122BOX, ThermoFisher, Waltham, MA, USA) which were run at 200V for 45 minutes.

### Immunocytochemistry

Neurons were rinsed briefly in warm (37°C) trisbuffered saline (TBS pH 7.4, diluted from 10X stock, 351-086, Quality Biological, Gaithersburg, MD, USA), then fixed for 5 minutes in a pre-warmed (37°C) solution of 4% paraformaldehyde (15710, Electron Microscopy Sciences, Morgantown, PA, USA) / 4% sucrose (S0389, Millipore Sigma, St. Louis, MO, UAS) / 1 mM EGTA (E14100-50.0, Research Products International, Mount Prospect, IL, USA) / 2 mM magnesium chloride (351-033-721, Quality Biological, Gaithersburg, MD, USA) in TBS, followed by 2 minutes in methanol at -20 °C. Fixation method was critical for signal quantification as longer fixation in paraformaldehyde (10-15 minutes) significantly decreased the endogenous phospho-tau signal. For surface GluA2 experiments, neurons were incubated with the GluA2 antibody in warm Neurobasal media (21103049, Gibco ThermoFisher, Waltham, MA, USA) for 10 minutes at 37°C prior to fixation. Immunostaining after fixation was performed as previously(Vossel et al., 2015) using TBS to replace phosphate-buffered saline in all solutions. Blocking was performed in 10% normal goat serum (S-1000-20, Vector Laboratories, Newark, CA, USA) in 0.01% triton X-100 in TBS, with primary and secondary antibodies incubated in 3% normal goat serum in 0.01% Triton X-100 in TBS. Coverslips were mounted using PermaFluor Aqueous Mounting Medium (TA-006-FM, Epredia, Kalamazoo, MI, USA) and maintained at 4°C until imaging. Imaging was performed on a Zeiss LSM 880 confocal microscope (Zeiss, Oberkochen, Germany). All image analysis and quantification were performed using ImageJ/Fiji. Dendrites and synapses were quantified manually, while blinded to treatment, and whole cell measurements were made using a cell mask based on the cell fill signal.

### ExRai-AKAR2 Measurements

Neurons were transfected and analyzed as previously(Zhang et al., 2021). Neurons were treated for 22-25 hours with 100 µM picrotoxin (1128, Tocris biosciences, Bristol, UK). Bicuculline was not used due to its fluorescence interfering with quantification in the 405 wavelength during live imaging. Excitatory (spiny) neurons were live-imaged 2 days after transfection in clear media on a Zeiss spinning-disk confocal microscope. Dual GFP excitation-ratio imaging was performed using the 488- and 405-nm lasers over a 20-minute period. The ratio of 488 to 405 signal was calculated from the maximum projected image and quantified as an average over the 20-minute imaging period for each neuron. All image analysis and quantification were performed using ImageJ/Fiji.

### Plasmid Cloning

GFP and tau plasmids were cloned into the pFSW backbone to limit transgene expression to neurons. Azurite was expressed using a pCAG backbone as published(Fang et al., 2021). The tau sequence (human 0N4R) was used from the pRK5-EGFP-Tau construct (#46904, Addgene, Watertown, MA, USA) with a myc tag added in frame to the N-terminus of the protein. Tau mutations were performed using primers designed with the assistance of NEBase Changer using GCA to encode alanine and the Q5 site-directed mutagenesis kit (E0554S, New England Biolabs, Ipswich, MA, USA). All inserts were confirmed with sequencing studies.

### Mice

Male and female C57/B6 wildtype mice were treated at 10-12 weeks of age with daily intraperitoneal injection of 1 mg/kg picrotoxin (1128, Tocris biosciences, Bristol, UK) freshly diluted in 2% DMSO in phosphate buffered saline (PBS) (2 day treatment n=3, 14 day treatment n=5), or a vehicle control (n=7). 24 hours after the final treatment, mice underwent deep anesthesia with transcardial perfusion of PBS followed by 4% paraformaldehyde. Brains were post-fixed in 4% paraformaldehyde for 24 hours and cut at 40µm thickness. All experimental procedures involving animals were approved by the Animal Care and Use Committee at Johns Hopkins University School of Medicine.

### Immunohistochemistry

Floating slices were rinsed 3 times in tris-buffered saline, then incubated 20 minutes at room temperature in LAB solution (#24310, Polysciences, Warrington, PA, UA). Streptavidin and biotin blocking was performed using the Streptavidin/Biotin blocking kit (SP-2002, Vector Laboratories, Newark, CA, UAS) then a mouse-on-mouse staining protocol was used to develop the phospho-tau signal using the M.O.M. immunodetection kit (FMK-2201, Vector Laboratories, Newark, CA, UAS).

Fluorescein signal was developed for 10 minutes prior to washing, then application of fluorescent secondary antibodies for the MAP2 and cfos signal. Slices were mounted on Superfrost Plus slides (22-037-246, Fisher Scientific, Hampton, NH) and coverslipped using VECTASHIELD (H-1000-10, Vector Laboratories, Newark, CA, USA). MAP2 signal was not quantified but used to ensure measured tau signal was in neuronal cell bodies. Brain slices from tau knockout mice(Dawson et al., 2001) were used during method development to ensure signal specificity given the low signal intensity of phospho-tau in wildtype mice.

### Experimental Design and Statistical Analysis

Neuronal culture experiments were performed in a minimum of 2 independent cultures and vehicle controls were included in all cultures. The number of neurons analyzed in each condition is listed in the figures. All imaging and analysis were performed blinded to the experimental treatment in culture and brain slices. Fluorescence intensity in each experiment was normalized to the average signal of the control neurons for that experiment to allow the data from independent experiments to be combined. For mouse brain imaging, signal intensity was normalized to the average signal in the outer molecular layer of the dentate gyrus in each slice, which did not differ between groups. Mouse brain data were normalized to the average control signal per experimental batch for graphing purposes, but not for statistical analysis in which batch was included as an experimental variable.

Statistical analysis of normally distributed data used t-test or one-way ANOVA with post-hoc testing, as in the results. Non-parametric equivalents were used with non-normally distributed data. Simple linear regression was used to test the association of cfos and pTau in neuronal culture. As multiple measurements were made within each brain slice for the mouse experiments, statistical testing was performed using mixed-effects linear models in R (version 4.5.1)(R Core Team, 2022)/R Studio (Posit Software, Boston, MA, USA) using the lme4 package (version 1.1-37)(Bates, 2015). Mouse was always used as a random intercept term to account for interindividual variability and use of multiple measurements in each mouse. pTau was modeled as an interaction between treatment and hippocampal region with batch treatment day as covariates. As treatment duration had no effect, it was removed from subsequent analyses. pTau was also modeled as an interaction between cfos and treatment with batch as a co-variate. To determine the regional effects, pTau in each treatment group was modeled separately as an interaction between hippocampal region and cfos with batch as a covariate. All data are available upon reasonable request to the corresponding author.

## Results

### Increased Endogenous Dendritic Tau Phosphorylation in Excitatory Pyramidal Neurons with Hyperexcitation

To examine the interaction between tau and neuronal activity, we quantified phosphorylated and total tau in primary rat hippocampal neurons with chronic changes in neuronal activity. Hippocampal neurons were treated with either 1µM tetrodotoxin (TTX), 20µM bicuculline, or a vehicle control for 24 hours. With bicuculline treatment, there was a small but significant increase in tau phosphorylation at the AT8 site (pS202/T205) by western blot with no change in total tau levels (Fig. S1A-C, One-way ANOVA with Dunnet’s *post hoc test*, df=32, *p=*0.03). While the change was subtle on western blotting, immunofluorescence showed a dramatic induction of endogenous tau phosphorylation in dendrites with chronic increased activity (Fig. 1A). This induction of tau phosphorylation occurred only in a subset of hippocampal neurons, accounting for the more subtle change in western blotting.

**Figure 1:**
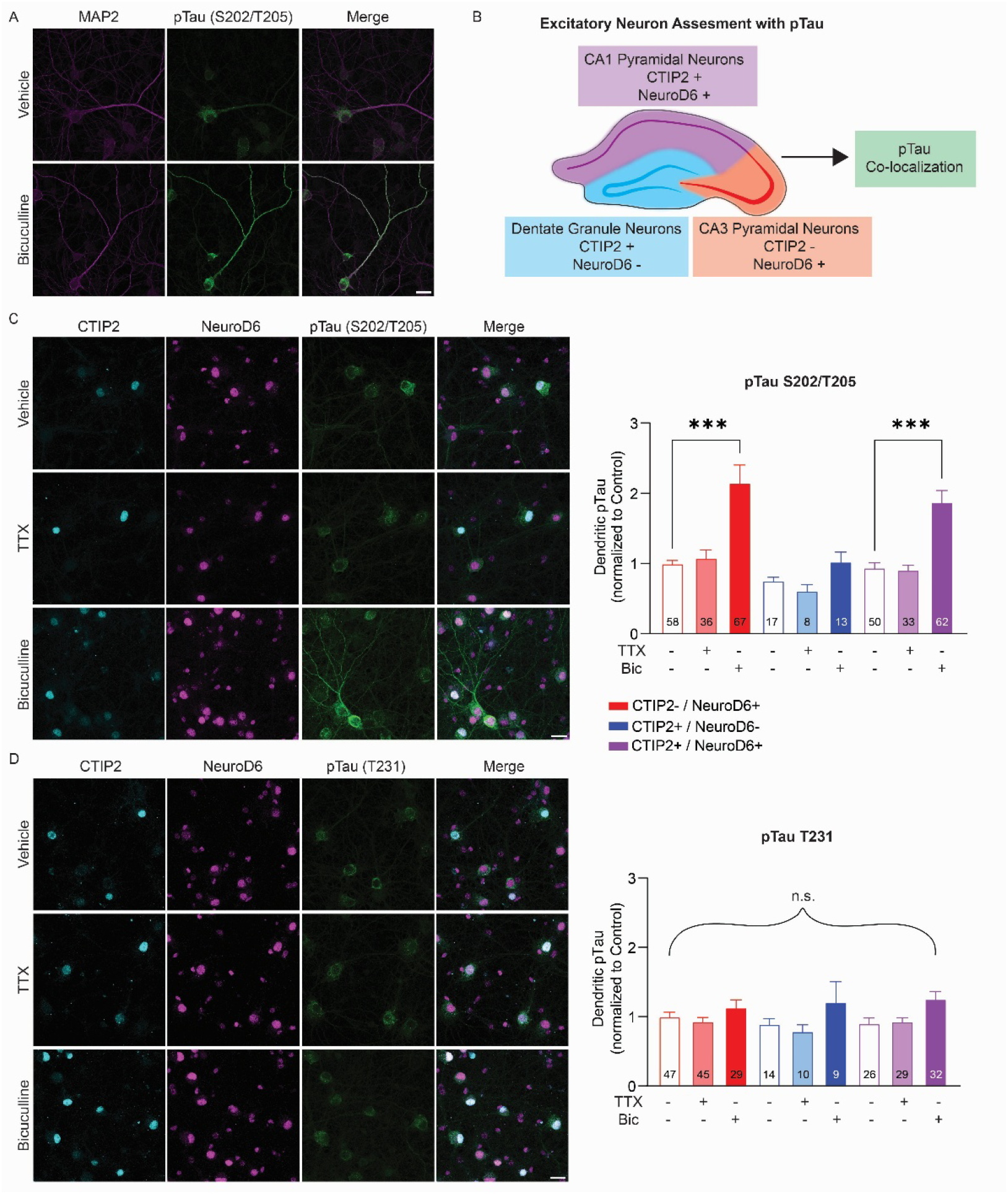
Endogenous Site-Specific Dendritic pS202/T205 Tau Phosphorylation in Excitatory Pyramidal Neurons with Increased Neuronal Activity. (A) Increased tau phosphorylation in dendrites (MAP2+) with 20 µm bicuculline treatment. (B) Schema of hippocampal excitatory neuron subtyping for colocalization with phospho-tau. (C) Increased tau phosphorylation at S202/T205 is induced by bicuculline specifically in excitatory pyramidal neurons (NeuroD6+) but not altered by TTX. (D) pT231 tau phosphorylation is not altered by changes in neuronal activity. Bic, bicuculline; TTX, tetrodotoxin. Scale bar is 25 µm. Neurons quantified in each condition are indicated in

To determine if the tau phosphorylation changes were affecting a specific neuronal population, we repeated the experiment with markers of specific hippocampal neuronal subtypes. To determine which population of excitatory neurons were involved, we took advantage of the differential expression of transcription factors in hippocampal excitatory neurons. Excitatory CA region pyramidal neurons in the hippocampus express NeuroD6(Bormuth et al., 2013), while dentate granule neurons and CA1 pyramidal neurons express CTIP2 (aka. *BCL-11B*)(Lee et al., 2013). By co-labeling with NeuroD6, CTIP2, and phospho-tau, we were able to differentiate putative CA1 pyramidal neurons (NeuroD6+, CTIP2+), CA3 pyramidal neurons (NeuroD6+, CTIP2-), and dentate granule cell neurons (NeuroD6-, CTIP2+) (Fig. 1B).

At baseline, dendritic tau phosphorylation at pS202/T205 was low and similar between all excitatory neuron types. Treatment with tetrodotoxin to decrease neuronal activity did not change dendritic tau phosphorylation. However, neurons treated with bicuculline showed increased dendritic tau phosphorylation specifically in CA1 and CA3 excitatory pyramidal neurons (Fig. 1C, Kruskal-Wallis with Dunn’s *post hoc* test, bicuculline vs. control CA1 and CA3 *p*<0.001). Dentate granule neurons showed no change in tau phosphorylation. The dendritic tau phosphorylation was also specific to the pS202/T205 AT8 site, as pT231 showed no alterations in any neuron type with tetrodotoxin or bicuculline treatment (Fig. 1D, Kruskal-Wallis with Dunn’s *post hoc* test, *p*=0.072). GAD67-positive cells had very low tau phosphorylation at baseline with no increase after bicuculline treatment (Fig. S1D), confirming that the neurons with dendritic tau phosphorylation were excitatory.

### Tau Phosphorylation is Associated with Neuronal Activity and Downstream of PKA

To begin examining mechanisms underlying this change in tau phosphorylation, we first examined the association with neuronal PKA inhibitor, KT5720, showed similar prevention of activation using cfos as a marker for neuronal activity(Chung, 2015). As expected, bicuculline treatment increased both tau phosphorylation and neuronal activation (Fig. 2A-C, Mann-Whitney test, pTau *p*=0.0009, cfos *p*<0.0001). There was a positive correlation between dendritic phospho-tau and cfos within neurons after bicuculline treatment (Fig. 2D, simple linear regression, bicuculline *p*=0.0062, R^2^=0.076), implying that tau phosphorylation within neurons was downstream of neuronal activation.

**Figure 2:**
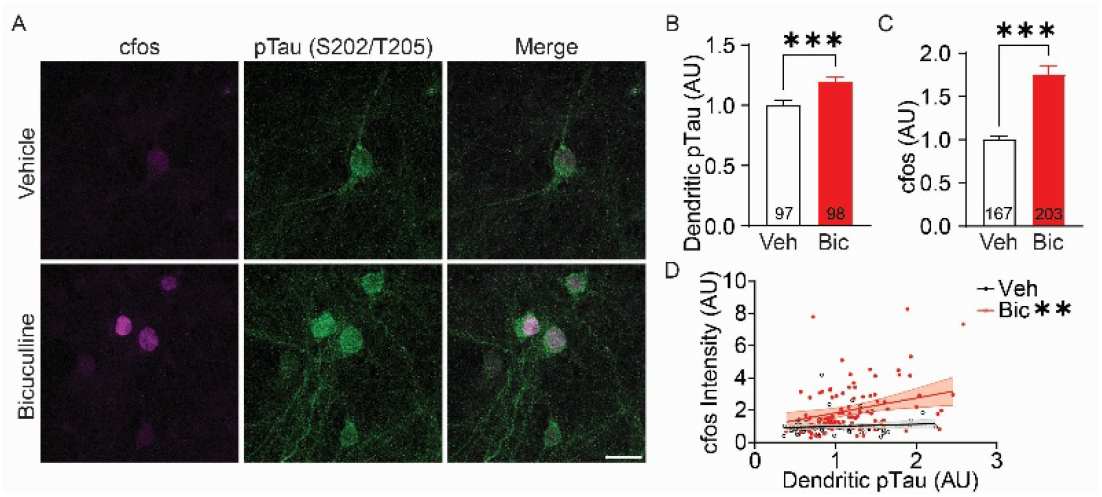
Endogenous Dendritic pS202/T205 Tau Phosphorylation Associated with Neuronal Activity. (A) Representative images showing higher dendritic tau phosphorylation in cfos-positive neurons with 20 µm bicuculline treatment. As expected, bicuculline treatment increased both (B) dendritic tau phosphorylation at S202/T205 and (C) cfos expression. (D) There was a direct correlation between dendritic tau phosphorylation and cfos expression in neurons with bicuculline treatment. Bic, bicuculline. Scale bar is 25 µm. Neurons quantified in each condition are indicated in graph. Error bars are SEM; shading is 95% CI. *, *p*<0.05; \*\**p<*0.01.

A targeted kinase screen was performed to determine which kinase was involved in endogenous dendritic tau phosphorylation during chronic neuronal activity (Table S1). Surprisingly, only PKA inhibitors prevented the increase in tau phosphorylation induced by chronic neuronal hyperexcitation (Fig. 3A-D). Pre-treatment with 5 µM H89, a PKA inhibitor, prevented the tau phosphorylation increase by bicuculline treatment in excitatory pyramidal neurons (NeuroD6+) (Fig. 3A,C, Kruskal-Wallis with Dunn’s *post hoc* test, bicuculline vs. control *p*<0.001, bicuculline vs. H89/bicuculline *p*=0.28). A second tau phosphorylation with increased neuronal activation (Fig. 3B,D, Kruskal-Wallis with Dunn’s *post hoc* test, control vs. bicuculline *p*=0.052, bicuculline vs. bicuculline/KT5270 *p*=0.0021, control vs. KT5270 *p*=0.021). Consistent with these findings, PKA activity was itself increased in dendrites with chronic increased neuronal activity (Fig. 3E,F, Mann-Whitney test, *p*<0.0001).

**Figure 3:**
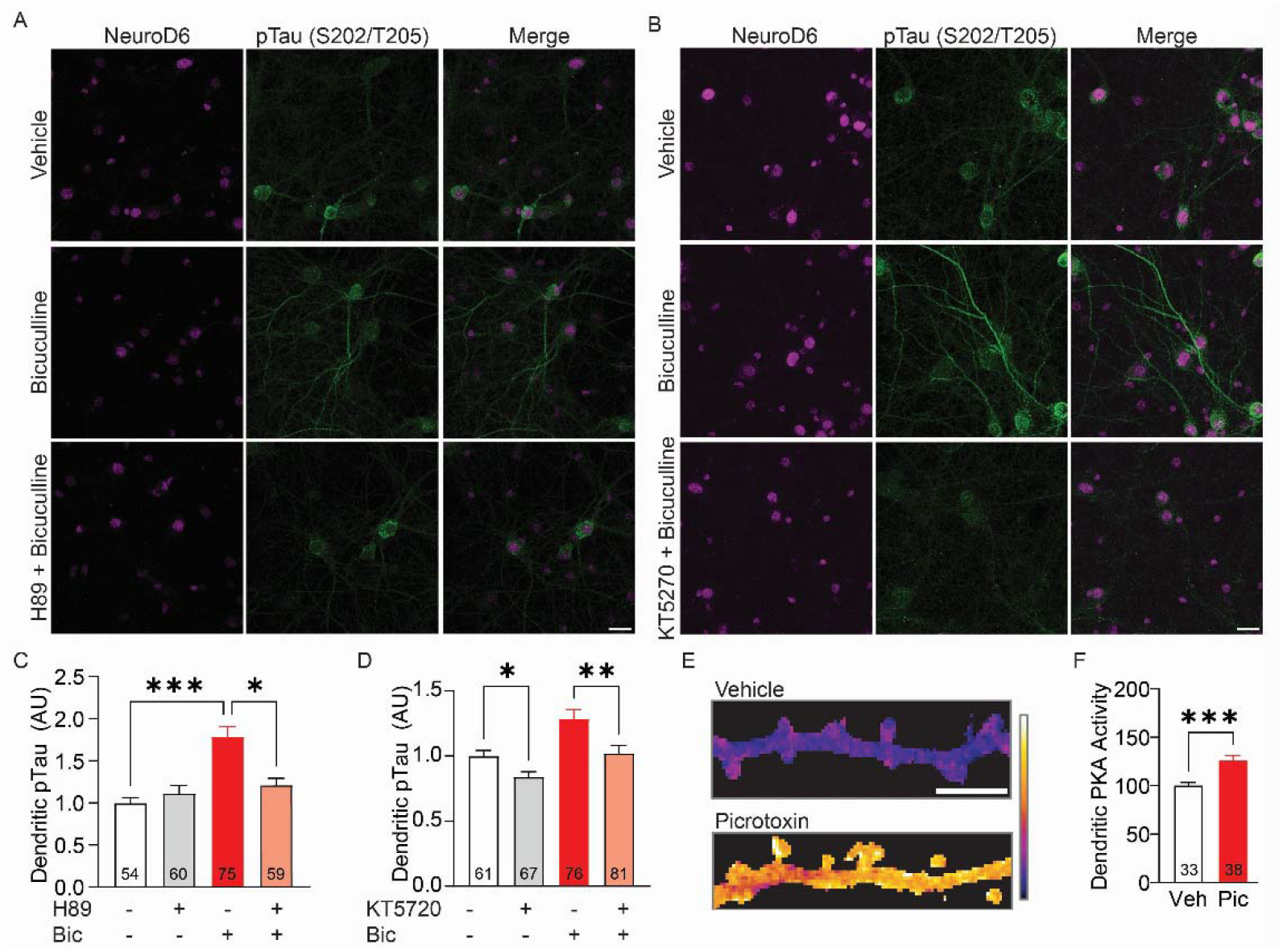
Activity-Related Endogenous Tau Phosphorylation is Downstream of PKA Activation. (A,B) Representative images of neurons were pre-treated with a PKA inhibitor, either 5 µm of H89 or 3 µm KT5720, for 10 minutes prior to bicuculline treatment. (C) H89 prevents increased dendritic tau phosphorylation with bicuculline treatment. (D) KT5270 pre-treatment prevents dendritic tau phosphorylation. (E,F) PKA activity was measured using ExRai-AKAR2 after 24 hours of 100 µM picrotoxin treatment. (E) Representative measurements of increased dendritic PKA activity in spiny excitatory neurons with increased neuronal activity and (F) quantification. Bic, bicuculline; Pic, picrotoxin; Veh, vehicle. Scale bars are 25 µm (A,B) and 5 µm (E). Neurons quantified in each condition are indicated in graph. Error bars are SEM. *, *p*<0.05; \*\**p<*0.01; ***, *p*<0.001.

### Tau Phosphorylation Regulates Surface GluA2 Expression

Tau knockout and pseudo-phosphorylation experiments have shown a role for tau in modulating AMPA receptors during a variety of neuronal activity changes. Because tau knockout alters GluA2 trafficking and GluA2 regulation is critical to homeostatic scaling in neurons(Suzuki and Kimura, 2017; Yong et al., 2020), we asked whether modulating phosphorylation specifically at the AT8 site would alter GluA2 regulation. Based on prior literature, the AT8 antibody primarily detects dual phosphorylation at the pS202/T205 site, though it can recognize tri-phosphorylation including S199(Goedert et al., 1995; Porzig et al., 2007). These three amino acids were therefore mutated to alanine in a myctagged human tau construct (0N4R) to prevent AT8 phosphorylation (mycTau-AT8A: S199A/S202A/T205A). Overexpression of myctagged wildtype-tau in neurons showed a high level of AT8 phosphorylation, which was absent in the AT8A-mutant tau (Fig. 4A).

**Figure 4:**
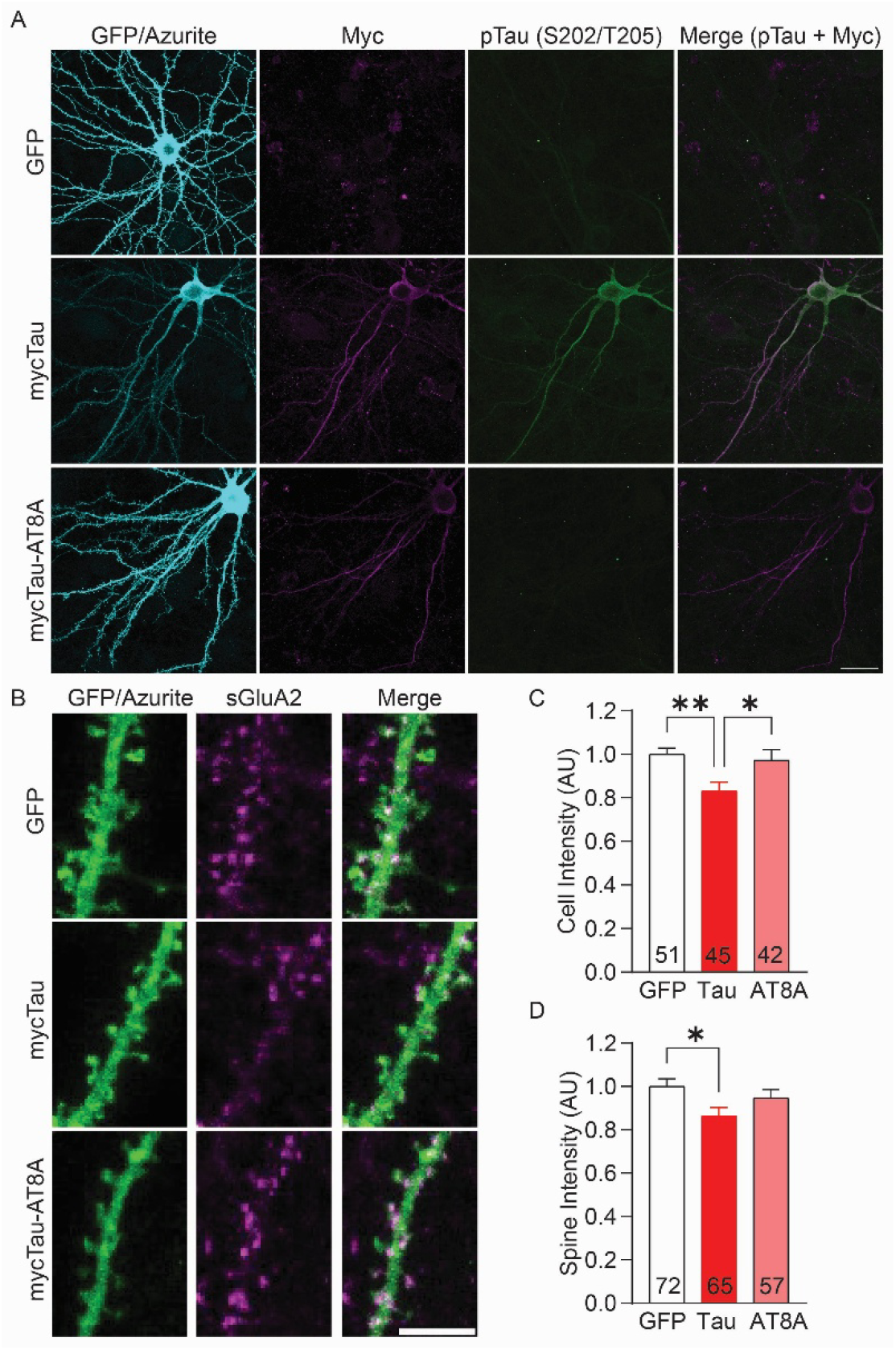
Tau Phosphorylation at the AT8 Site Modulates Surface GluA2 Expression. (A) Transfected hippocampal neurons showing expression of GFP, myc-tagged tau, and myc-tagged tau mutated at the AT8 sites with lack of AT8 staining. Neurons were co-transfected with azurite in all experiments. (B) Endogenous surface GluA2 levels in spines and dendrites of hippocampal neurons transfected with myc-tau, myc-tau with a phospho-mutant AT8 phosphorylation site, and GFP as a control. (C) Loss of surface GluA2 in neurons overexpressing tau, which is prevented by mutation of the AT8 phosphorylation site. (D) Loss of surface GluA2 in spines with overexpressing tau, with a trend to prevention by mutation of the AT8 phosphorylation site. Scale bar is 25 µm (A) and 5 µm(B). AU, arbitrary units normalized to control. Neurons quantified in each condition are indicated in graph. Error bars are SEM. \**p<*0.05; **, *p*<0.01.

Expression of wildtype tau decreased overall surface GluA2 levels in neurons, which was prevented in the phospho-mutant AT8A tau (Fig. 4B,C, One-way ANOVA with Tukey *post hoc* test, df=137, tau vs. GFP *p*=0.0053, tau vs. AT8A *p*=0.032). Similar reduction of surface GluA2 was seen with tau expression in post-synaptic spines, though the phospho-mutant tau only tended to improve post-synaptic surface GluA2 (Fig. 4D, Oneway ANOVA with Tukey *post hoc* test, df=193, tau vs. GFP *p*=0.023, tau vs. AT8A *p*=0.28), potentially suggesting tau may play a role in post-synaptic GluA2 trafficking outside of this specific phosphorylation site as well.

### Chronic Hyperexcitation Increases Endogenous Phospho-Tau in Mouse Hippocampus

While phospho-tau changes were selective and robust in hippocampal cultured neurons, we wanted to determine whether these changes also occurred *in* vivo. To examine endogenous tau phosphorylation in mice with chronically increased neuronal activity, we used chronic low-dose picrotoxin injection at sub-seizure treatment levels, which induces homeostatic down-scaling in CA3 pyramidal neurons(Lee et al., 2013). Mice were treated daily with intraperitoneal injection of 1 mg/kg picrotoxin for either 2 or 14 days, and sacrificed 24 hours after the last dose.

Low dose picrotoxin treatment increased S202/T205 phosphorylation tau in CA3 neurons in mice (Fig. 5A,B). 2 days of treatment were sufficient to see the effect in CA3 with no additional increase in phosphorylated tau after 14 days of treatment. *In vivo*, the effect on tau phosphorylation mainly localized to neuronal cell bodies. As the tissue was harvested 24 hours after the last injection, there were no changes in cfos expression with treatment (Fig. 5A) and there was no association between cfos expression and tau phosphorylation in any region in control animals (Fig. 5C). However, neurons with increased cfos expression were associated with higher levels of tau phosphorylation after picrotoxin treatment (Fig. 5C, mixed effects linear model, df=915, treatment by CA3 interaction *p*=0.0012). Similar to the findings in culture, this association was primarily driven by changes in CA1 and CA3 neurons (Fig. 5D, mixed effects linear model of picrotoxin-treated group, df=518, cfos:CA3 interaction *p*=0.0001, cfos:CA1 interaction *p*=0.0001).

**Figure 5:**
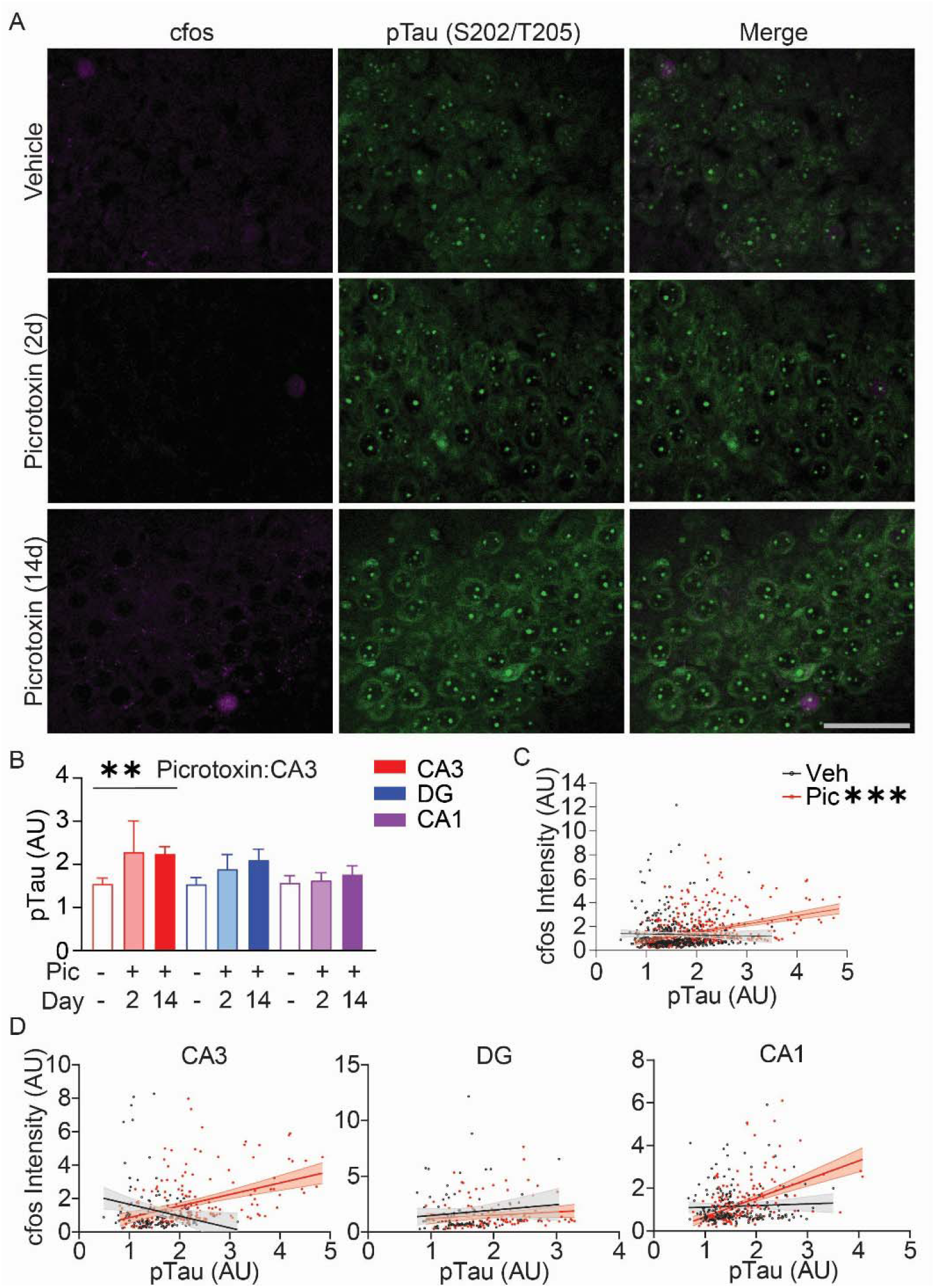
Endogenous pS202/T205 Tau Phosphorylation Associated with Neuronal Activity in Mouse Hippocampus. Mice were treated with 1mg/kg picrotoxin daily for 2 days (n=3) or 14 days (n=5), or treated with a vehicle control (n=7). (A) Representative images of CA3 region showing higher tau phosphorylation in the cell body of neurons with chronic low-dose picrotoxin treatment. The nuclear signal is non-specific staining as this was also present in tau knockout controls (not shown). (B) Cell body tau phosphorylation is increased at S202/T205 in CA3 by picrotoxin treatment. (C) Endogenous tau phosphorylation is increased in neurons with higher cfos expression with picrotoxin treatment (*p*<0.0001, treatment by cfos interaction, mixed effects linear model). (D) The positive association in picrotoxin treated mice was largely driven by CA3 and CA1 neurons. There was no association between cfos and phospho-tau with vehicle treatment (mixed effects linear model, df=391, *p=*0.14-0.41 by region). Statistics by mixed effects linear model. Pic, picrotoxin, Veh, vehicle. Scale bar is 50 µm. Error bars are SEM; shading is 95% CI. \*\**p<*0.01, \*\*\**p*<0.001.

## Discussion

This study identified selectively increased endogenous tau phosphorylation in CA1 and CA3 excitatory pyramidal neurons during chronic hyperexcitation. This tau regulation *in vitro* and *in vivo* was specific to excitatory pyramidal neurons, not occurring in dentate granule neurons or inhibitory neurons. Activity-related tau phosphorylation was also sitespecific, affecting the multiphosphorylation S202/T205 site but not the nearby pT231 site in the proline-rich region. Endogenous tau phosphorylation was directly associated with neuronal activity as measured by cfos expression. In cell culture, activity-related tau phosphorylation was downstream of PKA activity, which was induced by chronic hyperexcitation. Expression of tau in neurons with mutations to prevent phosphorylation at the S202/T205 site showed that this phosphorylation site is involved in regulating surface GluA2, consistent with prior literature and suggesting a role for tau phosphorylation in AMPA receptor down-regulation.

These findings provide insight into the endogenous, cell-type specific regulation of tau in neurons. While tau is ubiquitously expressed in neurons, genetic ablation of tau has a regional and cell type-specific effect on the balance of excitation and inhibition(Chang et al., 2021). Of the excitatory neurons examined in prior studies, tau had a significant role in the balance of excitation/inhibition in cortical pyramidal neurons but little effect on dentate granule neurons, somewhat similar to the findings in this study. We identified hyperactivity-related tau phosphorylation in excitatory hippocampal pyramidal neurons which was absent in granule cell neurons. While these changes were directly related to neuronal activity and PKA activation, the mechanism governing cell-type specificity remains unclear. Given the differential regulation of PKA across brain regions(Mucignat-Caretta and Caretta, 2020), it is possible that excitatory pyramidal neurons could have distinct regulation of PKA activity promoting tau phosphorylation.

PKA activity was increased by chronic neuronal hyperexcitation and mediated hyperactivity-related tau phosphorylation at the pS202/T205 site. While biochemical studies have shown that PKA does not phosphorylate the pS202/T205 site directly, PKA is consistently shown to facilitate tau phosphorylation at the pS202/T205 site by other kinases(Singh et al., 1995; Liu et al., 2006; Sengupta et al., 2006). No other kinase inhibitors screened, including inhibitors of GSK-3β, CK2, or CDK5, prevented the changes in tau phosphorylation. If PKA is priming the pS202/T205 site for phosphorylation, as indicated by prior literature, the other involved kinase is unknown. It is also possible that PKA is modulating local protein phosphatase activity(Leslie and Nairn, 2019), resulting in increased tau phosphorylation.

Tau phosphorylation has been implicated previously in AMPA receptor trafficking, consistent with this study. Tau has numerous post-translational modifications in normal brain, including regions of multiple clustered phosphorylation sites such as pS202/T205 which were once thought to exist only in a disease context(Morris et al., 2015a). Such abundant post-translational modification in normal brain suggests a crucial role for physiologic tau regulation in the brain which we are only beginning to understand. Tau knockout studies have established a role for tau in AMPA receptor trafficking during acute regulation of neuronal activity(Suzuki and Kimura, 2017; Prikas et al., 2022). However, most studies of tau phosphorylation have examined the relationship to AMPA receptors only in a disease context. One study of tau phosphorylation in P301L-mutant tau showed that phosphorylation in the proline rich region was critical for AMPA receptor downregulation by mutating 5 sites: S202, T205, T212, T217, and T231(Teravskis et al., 2020). The findings in this study refine the phosphorylation sites involved in AMPA receptor regulation, specifically implicating S202 and T205 which overlap between this study and the prior study. Our study also implicates tau phosphorylation in GluA2 regulation in a more homeostatic context, with increased expression and phosphorylation of wildtype tau sufficient to induce changes in surface GluA2, which are rescued by preventing phosphorylation at the AT8 site.

The neuronal cell-type specific regulation of tau phosphorylation may play a role in selective vulnerability in Alzheimer’s disease (AD), which shows early susceptibility of excitatory pyramidal neurons to tau pathology. Tau aggregates are thought to spread through the brain in AD via functional neuronal connections(Rudman et al., 2026). Given this hypothesis, tau pathology from entorhinal cortex should spread to both dentate gyrus and CA1 in human AD. However, dentate granule neurons are resistant to tau pathology until late in the disease course despite being highly connected to entorhinal cortex(Braak et al., 2006). It is tempting to speculate that activity-related tau phosphorylation could make neurons more vulnerable or resistant to tau pathology in AD. The combination of tau spread through functional neuronal connections and selective vulnerability associated with hyperexcitation-related tau phosphorylation in AD could potentially explain why CA1 accumulates early aggregates while dentate granule neurons are resistant. However, more studies would be needed to determine how and whether activity-related tau phosphorylation is related to vulnerability to tau aggregation in disease.

## Acknowledgements

This work was supported by an award from the National Institute on Aging (K08AG070053) and generous support from the Oak Hill Fund. The content is solely the responsibility of the authors and does not necessarily represent the official views of the National Institutes of Health.

