## Supplemental Figure 1, Table S1 for "Neuronal Hyperactivity Induces Tau Phosphorylation Selectively in Excitatory Pyramidal Neurons Which Contributes to GluA2 Regulation"

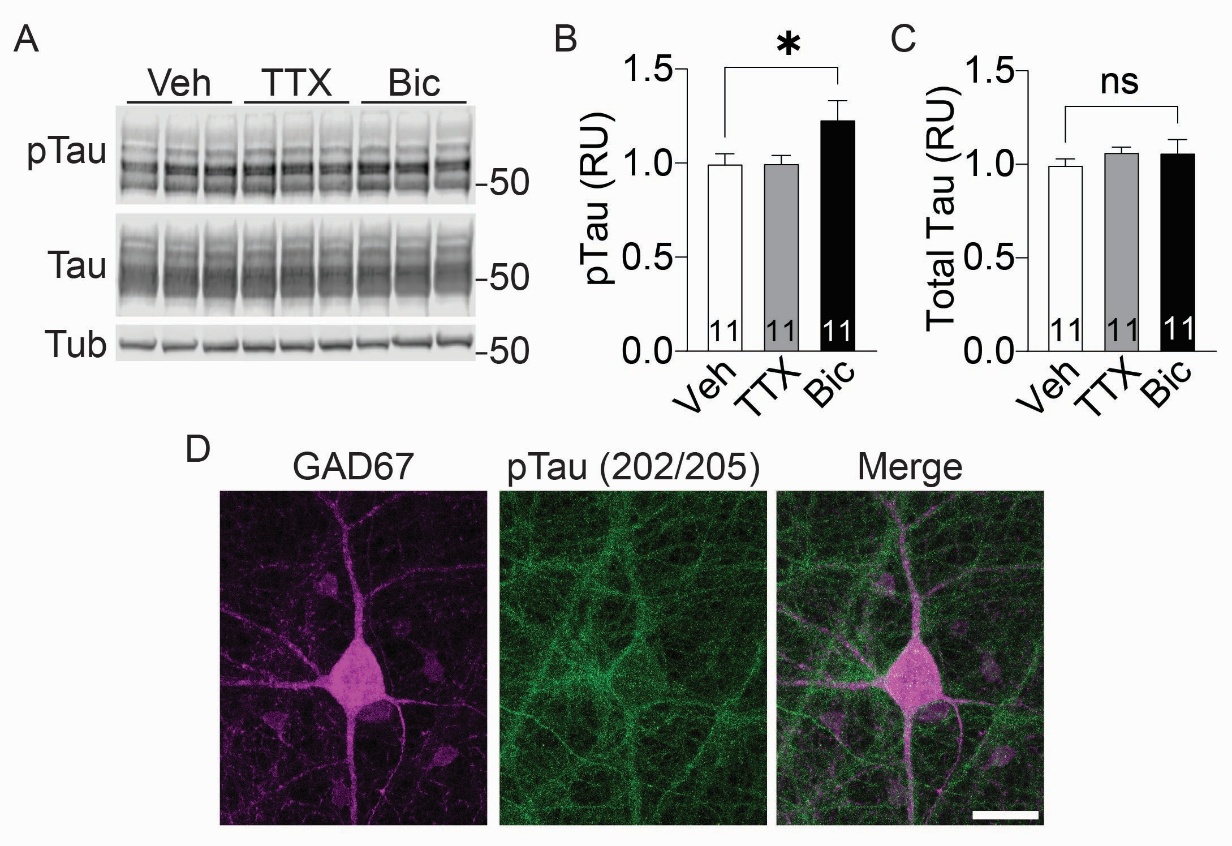

**Supplemental Figure 1: Tau Phosphorylation with Neuronal Activity Changes.** Rat hippocampal neurons were treated with TTX or bicuculline for 24 hours. (A) Western blot showing subtle increased tau phosphorylation (pS202/T205 normalized to total tau) with bicuculline treatment with (B) quantification. (C) Total tau levels (normalized to tubulin) did not change. (D) Representative image showing no increase in phospho-tau (green) in inhibitory interneurons (GAD67, magenta) with bicuculline treatment. Bic, bicuculline; RU, relative units; TTX, tetrodotoxin; Tub, tubulin; Veh, vehicle. Scale bar is 25 µm.

**Table S1: Kinase Inhibitor Screen**

| **Inhibitor** | **Target** | **Concentration** | **Dendritic pTau Effect** |
| --- | --- | --- | --- |
| Lithium chloride | GSK3β | 2 mM | No inhibition |
| GO6983 | PKC | 100 nM | No inhibition |
| (S)-Roscotivine | CDK5 | 20 µM | No inhibition |
| IC261 | CK1 | 20 µM | No inhibition |
| TBB | CK2 | 50 µM | No inhibition |
| SB203580 | P38 MAPK | 20 µM | No inhibition |
| Ly294002 | PI3K | 20 µM | No inhibition |
| U0126 | MEK1/2 | 10 µM | No inhibition |
| H89 | PKA | 5,10 µM | Inhibited |
| KT5720 | PKA | 3 µM | Inhibited |
